# Autosuggestion and Mental Imagery Bias the Perception of Social Emotions

**DOI:** 10.1101/2025.01.09.632121

**Authors:** Kasia A. Myga, Matthew R. Longo, Esther Kuehn, Elena Azañón

## Abstract

Cognitive processes that modulate social emotion perception are of pivotal interest for psychological and clinical research. Autosuggestion and mental imagery are two candidate processes for such a modulation, however, their precise effects on social emotion perception remain uncertain. Here, we investigated how autosuggestion and mental imagery, employed during an adaptation period, influence the subsequent perception of facial emotions, and to which extent. Separate cohorts of participants took part in five experiments, where they either mentally affirmed (autosuggested, Experiments 1a and 1b) or imagined (Experiment 2) that a neutral face would be expressing a specific emotion (happy or sad). Subsequent facial emotion perception was then assessed by calculating points of subjective equality (PSEs) along a happiness-sadness continuum. Our results show that both autosuggestion and mental imagery induce a bias toward perceiving facial emotions in the direction of the desired emotion, with larger Bayes factors supporting autosuggestion. Experiment 3 confirmed no effects when emotional words were presented instead, suggesting a reduced role of response bias to drive this effect. Finally, experiment 4 validated the experimental setup by demonstrating standard contrastive aftereffects when participants are adapted to actual, physical emotional faces. Together, our findings provide an initial step toward understanding the potential of intentional cognitive processes to modulate social emotions, specifically by biasing emotional face perception. With comparable effect sizes observed for both autosuggestion and mental imagery, both strategies show promise for self-directed interventions. Their practical applicability may vary due to individual responses, preferred cognitive strategies, and potential overlaps in underlying cognitive mechanisms.

## 1. Introduction

Cognitive processes that effectively modulate emotion perception are of pivotal interest for psychological and clinical research. There are different methodologies to modulate emotion perception, such as, for example contextual cues (Reschke & Walle, 2021), hypnosis (Zhang et al., 2024), or mental imagery (Diekhof et al., 2011). Autosuggestion and mental imagery can be applied most flexibly given that no external person is needed (as in suggestion and hypnosis), and no specific equipment/environment needs to be used (as in meditation). It has therefore been argued that more research is needed to explore the potential of autosuggestion and mental imagery to modulate clinically relevant states, such as emotions, or pain (e.g., Berna et al., 2012; Sari et al, 2017; Shilpa et al., 2020; Myga et al., 2022, 2024b).

Consequently, in our approach, we follow the definition of autosuggestion detailed in Myga et al., 2022. Specifically in this context autosuggestion reflects a process where participants internally repeat suggestions aimed at shaping their perception of emotions in another person. Imagery, on the other hand, is defined as the process of retrieving perceptual information from memory, leading to the re-creation of perceptual experiences in the absence of sensory input (Kosslyn et al., 2001) This process relies on four key abilities: image inspection, offline image generation, mental image transformation, and mental image retention (Kosslyn et al., 1995).

With respect to autosuggestion, little is known about the effect of the intentional and internal reiteration of thoughts (e.g., “she is happy/sad”) on the subsequent perception of emotions. Previous studies have reported the beneficial effects of reiterating thoughts related to desired health outcomes (Sari et al., 2017; Shilpa et al., 2020). For instance, acutely ill geriatric patients suffering from multipathological conditions reported a greater quality of life after a one-month autosuggestive procedure (Sari et al., 2017). These effects were accompanied by physiological changes, such as increased cortisol levels, reaching the norms observed in healthy adults. Other studies have shown that autosuggestion can reduce unhealthy snack consumption (Ludwig et al., 2014) and support the use of other mental techniques such as autogenic training, a psychophysiological form of therapeutic and relaxation technique (Schlamann et al., 2010).

The effect on emotion perception, however, was not the focus of this previous research, and such approaches have not been compared to the potential effects of mental imagery. Recently, we have shown that autosuggestion modulates the intensity of tactile perception at the hand (Myga et al., 2024b); however, this investigation did not focus on emotion perception but on neutral, non-emotional touches. Nevertheless, this study shows that autosuggestion can be used in a standard experimental setting to modulate participants’ perception dynamically, and hence highlights its potential applicability to also modulate emotion perception.

With respect to imagery, exposure to visual imagery created in the “inner eye” has been shown to influence subsequent perception of emotions. For instance, Zamuner and colleagues (2017) demonstrated contrastive aftereffects, in which prolonged imagery of a face displaying one of six basic emotional expressions (anger, disgust, fear, happiness, sadness, or surprise) led to subsequent test faces of the same emotion, but with weaker valence, being perceived as more neutral. These results were similar to those obtained with adaptation to real emotional faces, albeit weaker in magnitude.

Here, we investigate the modulation of emotion perception via both autosuggestion and imagery. This allows us to directly compare the effectiveness of both approaches to modulate emotions in desired directions, which may have important clinical implications. We focus on social emotion perception given that well-controlled experimental setups exist on emotion perception when observing faces (Butler et al., 2008; Sou & Xu, 2019; Webster et al., 2004; Yuan et al., 2024). In addition, the ability to modulate social emotion perception is an important skill and many clinical interventions in psychiatry target this function in patients (Saccaro et al., 2024). People with depression (Bourke et al., 2010) and those on the autism spectrum (Harms et al., 2010) for example display negative biases in their emotion recognition. Penton-Voak and colleagues (2013) used a modified feedback paradigm to experimentally bias the perception of ambiguous faces as happy rather than angry in healthy adults and adolescents at risk of a criminal offense. Participants were given feedback that faces that had been judged as angry actually expressed a happy emotion. After the procedure, participants in the “modified feedback group” reported lower degrees of anger in their face ratings as compared to participants in the “accurate feedback group”. Interestingly, within the same study, similar results were found in violent offenders serving time in prison (Penton-Voak et al., 2013). These findings underscore the potential of interventions that can bias social emotion perception.

In this study, we use an adaptation paradigm, wherein participants are briefly exposed to a facial expression (suggested or imagined), to investigate cognitive processes that modulate subsequent social emotion perception. Specifically, we examine the effects of autosuggestion (Experiments 1a and 1b) and mental imagery (Experiment 2) and compare these to the influence of emotionally charged words (Experiment 3). Additionally, we contrast these findings with the effects of direct exposure to actual emotional facial expressions (rather than imagined or suggested), known to produce contrastive aftereffects (Experiment 4, “perceptual aftereffect”; Rhodes et al., 2010). Our aims are to determine (1) if autosuggestion and mental imagery can be used by participants to bias emotional face perception, (2) whether this effect is driven by response bias, and (3) whether effect sizes are comparable to those of perceptual aftereffects.

More specifically, in Experiment 1a, participants were presented with a neutral female face and instructed to use autosuggestion to influence their perception of the neutral adaptor, repeating phrases such as “she is happy” or “she is sad”. Autosuggestion in this context is defined as the intentional generation and reiteration of thoughts to alter facial emotion perception (Myga et al., 2022, 2024b). They then rated the emotional valence of a subsequent emotional face ranging from very sad to very happy. In Experiment 1b, instructions were modified to minimize imagery use. Experiment 2 asked participants to imagine a previously seen neutral face as very happy or very sad after the face was no longer visible. This involved creating visual representations in the “mind’s eye” of the neutral face stored in short-term memory, transforming it to convey happiness or sadness in the absence of any physical image. Experiment 3 used “HAPPY” or “SAD” words as adaptors alongside a neutral face to control for cognitive expectations (Nichols & Maner, 2008; Orne, 1962), such as the belief that viewing happy faces predisposes individuals to perceive more happy faces. This experiment included the association between an emotion (in this case conveyed by an emotional word) and a neutral face in the adaptation phase, but did not involve a deliberate intention to perceive or mentally create a happy or sad face, as in Experiments 1a-b and 2. Finally, Experiment 4, involved direct exposure to extreme facial expressions to confirm the expected contrastive aftereffects (Ellamil et al., 2008; Pell & Richards, 2013; Ryu et al., 2008; Zamuner et al., 2017) and compare effect sizes across paradigms.

Together, we present a series of five experiments designed to investigate the effect of different cognitive strategies to modulate social emotion perception. We analyzed both the direction and strength of these effects while assessing the relative effectiveness of these techniques. The results of this study aim to identify potential applications for the modulation of social emotions to potentially transfer such techniques to clinical use.

## 2. Experiment 1a: Autosuggestion

Experiment 1a investigates whether autosuggestion biases the perceived emotional valence of an observed face in the direction of happiness or sadness. Specifically, participants were asked to perceive a neutral female face presented on the screen as happy or as sad while reiterating the respective thought (e.g., “the face looks happy”).

To determine the desired sample size, we based our calculations on the results of Zamuner and colleagues (2017). They compared aftereffects produced by perception, mental imagery, and baseline condition for six emotions: happy, sad, afraid, angry, disgusted, and surprised. We considered the effect size of the main effect of expression, which in their study was partial η^2^ = .359. We first converted their partial η^2^ effect size to Cohen’s d, which was equal to d*_z_* = .73. Next, we conducted a power analysis using G*Power version 3.1.9.7 (Faul et al., 2007). We set the alpha level to .05 and power to .8, which indicated a sample size of 17 participants was needed. However, we tested a few additional participants to account for the possibility of exclusions due to unforeseen issues, such as incomplete data or outliers.

### 2.1. Materials and methods

#### Participants

N = 23 participants were tested, from which n = 5 participants were excluded from the analyses. Exclusion was determined by the number of test trials in which participants were presented with one extreme of the stimulus spectrum – either the extremely happy morphed face (0) or the extremely sad morphed face (99) – for more than 50% of the trials in one staircase (see exclusion procedure below), compromising the intended variability in the test stimulus presentation. This occurred because some participants consistently indicated that the saddest face appeared closer to happy, and vice versa, leading to a strong bias in their responses. N = 18 participants formed the final sample (7 females, mean age = 25.56 years, SD = 2.99). All participants had normal or corrected-to-normal vision and were naïve to the purposes of the experiment. N = 16 participants were right-handed and n = 2 ambidextrous (mean = 86.08; range: 46.70 – 100; Edinburgh Handedness Inventory, Oldfield, 1971). Participants provided written informed consent for participation and were paid for their attendance. The procedures of this study were approved by the ethics committee at Otto-von-Guericke University Magdeburg, Germany (ethics number 01/19). Both the experimenter and the participants communicated in English. Each participant was tested individually.

#### Stimuli

Test stimuli were composed of two female facial expressions of a single person from the Karolinska Directed Emotional Faces (KDEF) database (Lundqvist et al., 1998) representing happy: AF29HAS and sad: AF29SAS emotions. These images were first cropped in a black oval mask (size 297 x 411 pixels) to remove hair using GNU Image Manipulation Program, GIMP version 2.10.32. Next, the images were converted into grayscale. The WinMorph 3.01 morphing tool was used to create a continuum of test stimuli transitioning in the range between happy (stimulus value 0) and sad (stimulus value 99). There were altogether 100 test stimuli created in the morphing step of one unit. Note, that the morphed levels corresponded to the relative mixture of the two stimuli, and not the perceived valence. In other words, a 49% morph represents an equal blend of the two faces, but may not be perceived as neutral or equally happy and sad. The perceived emotion depends on the criterion, the observer’s sensitivity, and the specific faces used to create the morphs. Test stimuli were the same in all five experiments. For an example of test stimuli, see **Figure 1** (a full set of stimuli can be found on Open Science Framework (https://osf.io/wk9xq/?view_only=aabe0f707ce243e180c1ed34a8d0bfc7). As an adapting stimulus, we used a neutral facial expression of the same identity selected from the KDEF database (Lundqvist et al., 1998): AF29NES (see **Figure 2b**). This image was processed the same way as the test stimuli (see above). The selection of the test stimulus on each trial was determined by a Bayesian adaptive algorithm QUEST (Watson and Pelli, 1983), based on the history of previous responses, and implemented using the Quest toolbox distributed as part of Psychtoolbox (Brainard, 1997; Kleiner et al., 2007).

**Figure 1.** Example of emotional facial expressions. Test stimuli #0, 12, 25, 37, 49, 62, 74, 86 and 99. Test stimuli were generated using the WinMorph 3.01 morphing tool, transitioning between a happy face (Stimulus #0) and a sad face (Stimulus #99), in the steps of one unit. The morphed levels represent the relative mixture of the two original stimuli. It is important to note that these levels correspond to the proportion of each original face in the blend, rather than the perceived emotional valence. A 49% morph is an equal blend of the happy and sad faces but may not necessarily be perceived as neutral or equally happy and sad. **\*Due to repository submission guidelines, the figure displaying facial stimuli was omitted from the article. To view the figure, please visit:** https://osf.io/wk9xq/?view_only=aabe0f707ce243e180c1ed34a8d0bfc7

**Figure 2.** **a) Trial sequence and adapting stimuli used in all experiments.** The experimental design was consistent across experiments encompassing two tasks in each trial: task 1 - exposure to the adaptor and task 2 - judgment of emotions in test faces. Note: in Experiment 2 (imagery), each trial started with the presentation of a neutral face for 1 s to remind participants of the identity of the person they were asked to keep in their minds, followed by a mask covering the full screen (see content of dotted blue line). Each trial continued (or started for the rest of the experiments) with the display of an adapting stimulus for 6 s (see **2b** for the specific adapting stimuli). Next, a green cross appeared on the screen indicating the end of the first task and the initiation of the second task. Next, a test stimulus appeared, and participants were asked to judge whether the test faces looked closer to the emotion happy or closer to the emotion sad. The face stimuli to judge were morphed faces of happy and sad expressions of the same female identity. The selection of test stimuli was determined by a Bayesian adaptive algorithm QUEST (Watson and Pelli, 1983), based on the history of previous responses. After a response was made, a white fixation appeared, to signal the end of the trial. **b)** During the adaptation phase in Experiment 1a-b (autosuggestion), participants were exposed to a neutral face and their task was to reiterate thoughts stating that the person they were observing looked very happy or very sad. To strengthen the effects of autosuggestion, they were instructed to perceive the noisy pixels on the image as contributing to the perception of the person’s happiness or sadness (Experiment 1a). In Experiment 1b this instruction was omitted and instead, participants were asked to avoid using imagery to their best abilities. In Experiment 2 (imagery), an oval shape with a blurred contour of the face was presented on the screen, and participants were asked to imagine the earlier seen neutral female face as happy or sad. In Experiment 3 (words), participants looked at the adapting stimuli containing the same neutral faces as in Experiment 1, however this time a word: “HAPPY” or “SAD” was presented beneath the face, corresponding to each experimental condition. In Experiment 4 (perception of physical stimuli), participants viewed an extremely happy or sad face, depending on the experimental condition. **\*Due to repository submission guidelines, the figure displaying facial stimuli was omitted from the article. To view the figure, please visit:** https://osf.io/wk9xq/?view_only=aabe0f707ce243e180c1ed34a8d0bfc7

#### Experimental procedure

Participants were first presented with an information sheet to familiarize them with the concept of autosuggestion and with the trial structure of the experiment (see: https://osf.io/wk9xq/?view_only=aabe0f707ce243e180c1ed34a8d0bfc7). They were instructed to generate and reiterate thoughts affirming that the face presented on the screen appeared either happy or sad, depending on the experimental condition. In addition, to potentiate the process of autosuggestion, they were encouraged to interpret ambiguous visual elements on the screen, particularly around the mouth and eyes, as indicative of the targeted emotional expression. Specifically, they were instructed to notice that the mouth corners are in fact pointing upwards or that the formation of wrinkles around the eyes indicates a happy emotion, while mentally repeating the sentence “she is happy”. In addition, participants received verbal guidance and engaged in exercises under the supervision of the experimenter. To train on the autosuggestion task, participants looked at the neutral facial expression and tried to autosuggest that this face in fact looks happy or sad, respectively. Subsequently, participants underwent comprehensive training for the entire experimental task. Specifically, they were shown the adapting and test stimuli and they were instructed which ones to autosuggest, and which ones to judge as happy or sad. This training was a shortened version of the actual experiment. The experimenter emphasized the importance of avoiding visual imagery or mental pictures during the procedure, explaining that imagery is a separate mental process that could influence the results. All participants confirmed their understanding of this instruction.

The main experiment consists of two conditions, labeled after the elicited emotion perception: happy and sad, with the order counterbalanced across participants, separated by a short untimed break in the middle. The only difference between the conditions was which emotion was autosuggested. Participants were seated approximately 56 cm from the screen. The stimuli were shown on a 24.4-inch monitor (resolution: 1920 x 1080 pixels; refresh rate: 59.940 Hz), on a black background. Stimuli were presented at a visual angle of approximately 17.26 degrees (height) and were centered on the screen. Stimuli were presented using Psychtoolbox (Brainard, 1997; Kleiner et al., 2007), running on MATLAB (Mathworks, Natick, MA).

Figure 2 illustrates an experimental trial. Participants were informed that each trial involved two tasks: autosuggestion and the judgment of the test stimuli. In the first task, participants were instructed to autosuggest that the face looked very happy or very sad, respectively. The second task required them to indicate whether the subsequently seen face looked closer to happy or closer to sad. Each trial began with a blank screen for 0.5 s. Next, a neutral face was shown for 6 s. This was succeeded by a green cross for 1 s, indicating the start of task 2 and the subsequent 0.5 s presentation of the test face stimulus. Each new test stimulus was selected by QUEST adaptive psychometric procedure (Watson & Pelli, 1983), from 100 possible test faces (morphed faces of happy and sad expressions of the same female face, see above). QUEST adaptive psychometric procedure adjusts stimulus presentation based on participants’ responses to estimate a threshold level in this case of 75%. The screen remained blank until the participant responded by pressing the left or right arrows, for sad and happy, respectively. Once a response was given, a white fixation cross appeared, indicating the end of the trial and the start of the next trial. Participants were told that the judgments of the test stimuli were an independent task from the autosuggestion task, and that the judgments given for each face should not be influenced by the perceptual outcome in the first part of the trial. Note that the term “long-staying face” rather than “adaptor” was used when explaining the task, in this and the following experiments. This phrasing was chosen to avoid biasing participants with expectations about the effects being studied.

There were 60 test trials in each condition, split into two QUEST staircases of 30 trials each, one initially increasing and the other decreasing. Each staircase was initiated with an extreme facial expression, either the extremely happy morphed face (0) or the extremely sad morphed face (99), to counteract possible response biases from the starting direction of the staircase. Each staircase was randomly interleaved on a trial-by-trial basis. At the end of each block, the point of subjective equality (PSE, i.e., the stimulus for which the participant was equally likely to judge it as happy or sad 50% of the time) for each staircase was calculated using a Bayesian adaptive algorithm QUEST (with the function QuestMean; Watson & Pelli, 1983). The PSEs from the two staircases were then averaged to derive a single estimate for each condition, separately for the happy and sad conditions. Participants underwent one self-paced, untimed break in the middle of each block. The experiment started with an initial 60 s exposure to the neutral face, and participants were asked to autosuggest, as described above. This long adaptation period was repeated after the middle pause. At the end of all experiments, participants were debriefed on the purpose of maintaining a continuous focus on the adapting stimuli.

Before and after the experiment, participants rated their mood on a visual analog scale (VAS), with 0 corresponding to feeling sad and 100 corresponding to feeling happy. Evidence suggests that one’s own moods may interact with emotion perception in others (Sel et al., 2015). At the end of the experiment, participants filled out the following questionnaires: a) the Spontaneous Use of Imagery Scale (SUIS; Kosslyn et al., 1998, 12-60 points) to assess the spontaneous use of visual imagery in everyday life, b) the Short Suggestibility Scale (SSS, 21-105 points), a condensed version of Multidimensional Iowa Suggestibility Scale (MISS)(Kotov et al., 2004) to assess participants’ overall compliance with suggestions, and c) the Vividness of Visual Imagery Questionnaire 2 (VVIQ2, Marks, 1973, 16-80 points) with eyes open and closed, to evaluate vividness of participants’ visual images. In addition, participants answered several open questions to gain insight into how participants performed autosuggestion. The questions, responses, questionnaire ratings and qualitative results are available in the Supplementary Materials on the OSF platform (https://osf.io/wk9xq/?view_only=aabe0f707ce243e180c1ed34a8d0bfc7).

### 2.2. Analyses

Points of subjective equality (PSEs) were calculated using a Bayesian adaptive algorithm QUEST (Watson & Pelli, 1983), which estimated the percentage of morph at which participants were equally likely to judge the test face as happy or as sad. We obtained two PSEs per condition, which were averaged, resulting in one PSE for each participant and condition.

We used a paired sample t-test to compare the two PSE values between positive and negative conditions, with the significance level set to .05, and reported Cohen’s d*_z_* as a measure of effect size. When the assumption of normality was violated (Experiments 1a-b and 2), we conducted the non-parametric Wilcoxon signed-rank test in addition, and report the results of both tests. Bayes factors (*BF*10) were calculated to quantify evidence in favor of the alternative hypothesis compared to the null hypothesis (using a default Cauchy prior of 0.707 in JASP version 0.19.1.00).

#### Exclusion Criteria

Participants who were exposed to extreme test stimuli – either the extremely happy (0) or sad (99) morphed faces – for more than 50% of trials in a staircase were excluded, as this pattern indicated a strong response bias that compromised threshold estimation in the QUEST adaptive psychometric procedure. N = 5 participants were excluded based on this criterion. This exclusion criterion also addressed additional issues in some of these participants, including thresholds exceeding the 0-99 range or significant discrepancies between thresholds obtained across the two staircases within the same condition. Among the included participants, all experienced extreme trials (either the happiest or saddest face) in 43% or fewer of their trials, with most encountering extreme stimuli only on the first trial.

### 2.3. Results

There was a significant difference between PSEs in the happy versus sad autosuggestion condition (see Figure 3a). More precisely, in the happy condition, the PSE was higher (i.e., a facial morph closer to sadness, M = 48.75, SD = 16.28), meaning participants required a facial morph closer to a sad expression before identifying it as sad. In contrast, in the sad condition, the PSE was lower (M = 37.61, SD = 16.07), indicating that participants judged faces as sad even when they displayed more pronounced happy features. This shift in PSE in the direction of the adaptor demonstrates an assimilative aftereffect. The effect was statistically significant (t(17) = 3.144, p = .006) with a medium to large effect size (d*_z_* = .741) and strong evidence supporting the alternative hypothesis (BF10 = 13.372). We report the Wilcoxon signed-rank test for completeness, as the assumption of normality was violated: z = 2.504, p = .01, r = .673.

**Figure 3.**
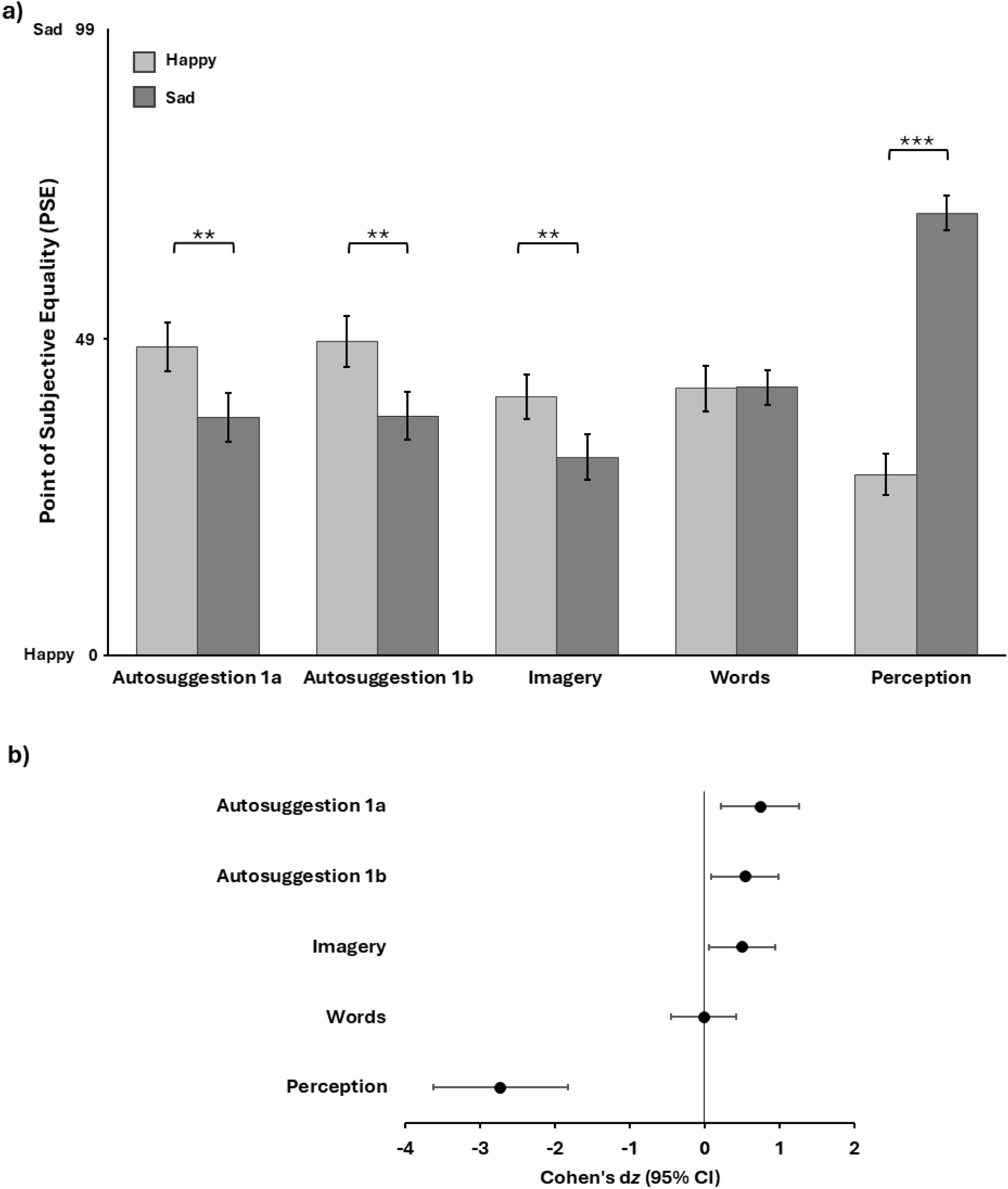
Overview of experimental results. **a) Comparison between PSE in happy and sad conditions in all experiments.** Both experiments on autosuggestion (Experiment 1a and 1b) as well as imagery (Experiment 2) produced assimilative aftereffects: the PSE in the happy condition was significantly higher than the PSE in the sad condition, as indicated by the paired sample t-test. After prolonged autosuggestion or imagery of a neutral face as “happy” or “sad”, they perceived a greater number of test faces in the direction of the autosuggested or imagined emotion. In an experiment where adapting neutral faces were paired with emotion describing the words: “HAPPY” or “SAD”, no PSE difference was observed between the two conditions. Conversely, in the perceptual experiment, using extremely happy and sad faces as adaptors, contrastive aftereffects were observed: the mean PSE in the happy condition was significantly lower than in the sad condition, indicating test faces were perceived in the opposite direction to the adapting emotion. Error bars represent Standard Error. Asterisks indicate significance level at p values: ** for p ≤ .01 and *** for p ≤ .001. **b) Effect size by experiments (Cohen’s dz).** The solid vertical line represents no effect of experimental manipulation. Each circle represents the point estimate of the effect size for the study corresponding to the label on the left. The lines on the left and the right of the circle represent the lower and higher 95% CIs for the effect size. Points to the left of the line represent contrastive adaptation aftereffects, and points to the right of the line indicate assimilative aftereffects.

For transparency, we repeated the analysis including the five outliers but excluded only the affected staircase data. Although this approach includes less robust data due to fewer trials (and the likelihood that participants with strong biases in one staircase may not provide accurate estimations), the results remained significant (PSE happy M = 47.57, SD = 17.73; PSE sad M = 39.62, SD = 16.46; t(22) = 2.285, p = .032, d*_z_* = .477; BF_10_ = 1.871).

#### Mood, suggestibility and imagery scores

Participants’ mood ratings at the start (M = 70.80, SD = 25.68) and the end (M = 64.83, SD = 26.33) of the experiment showed no significant difference (t(17) = 1.070, p = .300, d*_z_* = .252). Their mean suggestibility score (M = 49.94, SD = 9.91; Kotov et al., 2004), spontaneous imagery use (M = 41.17, SD = 7.49; Nelis et al., 2014, Sample 1), and vividness of imagery (combined eyes open/closed M = 67.08, SD = 10.53;) aligned with prior studies (Azañón et al., 2024; Tabi et al., 2022).

#### Task strategies overview from follow-up questionnaires

Participants were instructed to repeat autosuggestive sentences while interpreting ambiguous visual elements, particularly around the mouth and eyes, as indicating specific emotional expressions. Self-reports indicate that participants primarily focused on facial analysis for autosuggesting emotions rather than verbal suggestion, with 83% reporting imagery use overall. 83% of the participants reported changing their own expressions and mood shifts during the experiment, in alignment with the autosuggested emotion. A minority (11%) superimposed autosuggested emotions onto the screen, while 17% relied on verbal suggestions with complementary techniques. For a comprehensive overview of the self-reports and questions, please refer to the Supplementary Material (https://osf.io/wk9xq/?view_only=aabe0f707ce243e180c1ed34a8d0bfc7).

## 3. Experiment 1b

The results of the first experiment indicate a reliance on imagery and personal emotional engagement during the autosuggestion process with minimal use of repetitive verbal phrases (e.g., “this person is happy”, “this person is sad”). This made it difficult to assess whether the observed effects could also emerge when using suggestion in the form of intentional thought repetition. To address this, we conducted a second experiment with stricter instructions to further discourage participants from utilizing imagery during the autosuggestion task.

### 3.1. Materials and methods

#### Participants

Altogether N = 23 participants were tested, each seated in separate cabins. Data from one outlier was excluded based on the same exclusion criteria as for Experiment 1a, leaving n = 22 participants for analyses (12 females, mean age = 27.82, SD = 3.78). One participant was left-handed, one was ambidextrous, and the rest of the sample was right-handed (mean = 82.25, SD = 43.58, range: −100 – 100).

#### Stimuli

Stimuli were the same as those used in Experiment 1b.

#### Experimental procedure

In this adapted version of experiment 1a, we emphasized to participants in the instructions that they should avoid imagery. Specifically, we removed guidance to interpret ambiguous visual elements – e.g., around the mouth and eyes – as indicative of specific emotional expressions. For instance, in the happy condition, we eliminated instructions like “notice that the mouth corners are pointing upwards” or “wrinkles around the eyes indicate happiness” while mentally repeating “she is happy”. Additionally, we included more targeted questions about the autosuggestion process (see: https://osf.io/wk9xq/?view_only=aabe0f707ce243e180c1ed34a8d0bfc7 for the detailed questions). In addition, we introduced a controlled 5-minute break between the two conditions, replacing the participant-controlled break, and collected mood ratings before and after each condition to account for possible mood fluctuations following autosuggestion. Finally, to reduce testing time, we administered the VVIQ2 questionnaire only with eyes open, as participants would need their eyes open to track dynamic changes on the screen if they were to use imagery.

### 3.2. Results

The results of the second experiment parallel those of the first Experiment 1a. Specifically, the PSE in the happy condition (M = 49.58, SD = 18.86) was significantly higher than the PSE in the sad condition (M = 37.84, SD= 17.48), with a medium effect size (t(21) = 2.504, p = .021, d*_z_* = .534), and strong evidence supporting the alternative hypothesis (BF_10_ = 15.642). We report the Wilcoxon signed-rank test for completeness, as the assumption of normality was violated: z = 2.613, p = .007, r = .636.

#### Mood, suggestibility and imagery scores

Participants’ mood ratings before (M = 65.04, SD = 22.34) and after the experiment (M = 61.55, SD = 20.17) did not differ significantly (z = .087, p = .945, r = .022). Similarly, no significant difference was observed in mood ratings before (M = 65.83, SD = 22.50) and after the happy condition (M = 64.77, SD = 17.97, z = 1.185, p = .248, r = .239). Although participants’ mood at the end of the sad condition (M = 54.45, SD = 23.08) was lower than at the start of that condition (M = 63.39, SD = 21.08), this difference only showed a trend toward significance (*z* = 1.860, *p* = .065, *r* = .463). Participants demonstrated average levels of suggestibility (M = 51.27, SD = 12.41; Kotov et al., 2004), spontaneous imagery use (M = 39.73, SD = 8.55; Nelis et al., 2014, Sample 1), and vividness of imagery (M = 58.73, SD = 10.11; Azañón et al., 2024, Tabi et al., 2022).

#### Task strategies overview from follow-up questionnaires

The majority of participants (86%) repeated autosuggestive sentences describing intended emotions. Mental imagery was used by 59%, and 36% reported superimposing imagined expressions in at least one condition. Participants moderately changed their facial expressions (M = 2.77, SD = 1.45; scale 1–5) and leaned slightly toward verbal suggestions over imagery (M = 2.55, SD = 1.22; scale 1 = verbal, 5 = visual). Perceived autosuggestion effectiveness averaged as “moderately” for both conditions (mean happy = 3.1 (SD = 1.07), mean sad = 3.6 (SD = .99)). Detailed questions and responses are available in the Supplementary Material (“Results Exp. 1b Autosuggestion” on OSF, https://osf.io/wk9xq/?view_only=aabe0f707ce243e180c1ed34a8d0bfc7).

## 4. Experiment 2: Mental Imagery

In Experiment 2, participants viewed a neutral face and, after it disappeared, were instructed to imagine the person as happy or sad (see Figure 2a). This approach differs from the autosuggestion experiments in the sense that rather than *attempting to modify* the perceived expression of a continuously present physical stimulus through suggestion, participants instead focused on *creating a new emotional image* in their minds.

Using a previously seen object as a basis for imagery is a common approach, and aligns with Kosslyn’s (1995) mental imagery framework involving image inspection, image generation, transformation and retention, where they maintain the altered image until the test trial. We expected that the prolonged imagery of a happy or sad face would influence subsequent perception. However, the direction of this effect remained uncertain due to conflicting findings in prior research, finding both contrasting and assimilative aftereffects (D’Ascenzo et al., 2014; DeBruine et al., 2010; Korolkova, 2018; Palumbo et al., 2017; Zamuner et al., 2017).

### 4.1. Materials and methods

#### Participants

N = 24 participants were tested individually. N = 1 did not complete the experiment and n = 1 was excluded following the same exclusion criteria as in experiments 1a and 1b. Therefore, n = 22 formed the final sample (10 females, mean age = 26.77, SD = 5.34, 18 right-handed, mean 65.33, range: −100 – 100).

#### Stimuli

The same stimuli from Experiments 1a-1b were used in Experiment 2, with the addition of a blurred oval image presented during the adaptation phase. The blurred image was used to improve the generation and vividness of the mentally imagined adaptor face, as suggested by D’Ascenzo and colleagues (2014). The blur was superimposed on the neutral emotion female face using the blurring tool available in GIMP version 2.10.32, and had the same size and location as the previously seen neutral face and the test faces.

#### Experimental procedure and analyses

The experimental procedure was similar to Experiment 1. The main difference was that instead of asking people to reiterate a thought internally that would influence their emotion perception, participants were asked to imagine the previously seen neutral faces as happy or sad, while looking at a blurred outline of a neutral face (see Figure 2b**: Exp. 2. Imagery**).

At the start of each condition and after each break, participants viewed the neutral face for 5 s and were instructed to memorize it to later imagine it as happy or sad. Then, a noise mask covering the full screen was displayed for 1 s to minimize after-images (created in Matlab with imnoise using salt and pepper noise, density = 1). Next, a blurred face was shown for 60 s, during which participants were required to imagine the previously seen face as very happy or very sad. After this adaptation phase, a test face determined by QUEST was presented for 0.5 s, and participants judged whether it appeared closer to happy or sad. A white cross marked the end of each trial for 0.5 s. Subsequent trials followed a similar structure with shorter durations: neutral face for 1 s, noise mask for 1 s, and blurred face for 6 s as the adaptor. There were 60 trials per condition, and 120 trials in the whole experiment.

Before the start of the experiment, participants were shown a demonstration of the imagery procedure, as well as the trial structure, with only 10 trials per condition. Just like in Experiments 1a-b, it was stressed to participants to evaluate the test faces independently from the outcomes in the imagery phase. Participants filled out the same questionnaires as in Experiment 1a, and open-ended questions where the word “autosuggestion” was replaced with the word “imagery”. Analyses were identical to Experiments 1a-b.

### 4.2 Results

The results mirror those of experiments 1a and 1b (see Figure 3**: Imagery**): The PSE in the happy condition (M = 40.93; SD = 16.39) was significantly larger than in the sad condition (M = 31.35; SD = 17.16), with a medium effect size (t(21) = 2.334, p = .030, d*_z_* = .498), and moderate evidence supporting the alternative hypothesis (*BF*10 = 5.239), suggesting an assimilative aftereffect. Specifically, in the “imagine sad” condition, participants identified faces with about 30% sad content as equally happy or sad, and in the “imagine happy condition”, a greater amount of sad content was required for this categorization. We report the Wilcoxon signed-rank test for completeness, as the assumption of normality was violated: z = 2.354, p = .017, r = .573.

#### Mood, suggestibility and imagery scores

Participants’ mood ratings did not differ before (m = 69.86; SD = 22.40) and after (m = 61.29; SD = 20.24) the experiment took place, *t*(21) = 1.567, *p* = .132, *d_z_* = .334. Participants displayed average suggestibility levels (m = 47.59, SD = 11.35), spontaneous use of imagery (M = 39.91, SD = 7.10), and imagery (combined open/close eyes mean = 63.21, SD = 9.12, Azañón et al., 2024, Tabi et al., 2022).

#### Task strategies overview from follow-up questionnaires

In brief, participants used various strategies to imagine happy and sad faces beyond simply imagining the preceding neutral face as happy or sad. For happy faces, 23% referenced personal or others’ happy experiences, while 41% did not specify their process. For sad faces, 27% referred to personal or others’ sad experiences, and 32% focused solely on the perceptual outcome. In terms of imagery techniques, 36% placed the imagined person in a specific situation, and 23% superimposed the imagined image on the blurred outline presented on the screen, and 59% reported changing their own mood to match the imagined emotion. For a detailed overview of the self-report, refer to the Supplementary Materials: Table S6 on OSF (https://osf.io/wk9xq/?view_only=aabe0f707ce243e180c1ed34a8d0bfc7).

## 4. Experiment 3: Emotional words

The preceding results suggest that autosuggestion and mental imagery influence the perceived emotion of observed faces in the direction of the autosuggested or imagined emotion. Experiment 3 explored whether this effect could partly result from a response bias or cognitive expectations (Nichols & Maner, 2008; Orne, 1962), such as the belief that viewing sad faces predisposes individuals to perceive more sad faces. To examine this, we added the words “HAPPY” or “SAD” under the neutral face as the adapting stimulus, without instructing participants to evoke any specific emotion. If judgments were biased by these words, similar aftereffects to those in Experiments 1a-b and 2 would be expected. If not, it would suggest that response bias is not the primary factor driving the results observed in the preceding experiments.

### 4.1. Materials and methods

#### Participants

Altogether N=21 participants took part in the experiment. Participants were tested in groups of up to 4 people in separate cabins. There were no outliers based on the above-introduced criteria. The analyzed sample (10 females) had a mean age of 26.14 (SD = 3.44). All participants were right-handed (M = 91.78; range: 57.10 – 100).

#### Stimuli

Adapting stimuli were neutral faces as in experiments 1a and 1b, but in experiment 3, the following written English words were presented below the face: “HAPPY” or “SAD” (see Figure 2b**: Exp. 3. Words**). The words were presented centrally, 2.15 degrees of visual angle below the face in capital letters (Calibri 44), and with a letter height of 1.13 degrees of visual angle. Note, that the size of adapting and test faces were identical across experiments, and that all communication between the experimenter and participants was in English).

#### Experimental procedure and analyses

The experimental procedure and analyses were identical to experiment 1a, with the only exception that the adapting stimulus contained the written word: “HAPPY” or “SAD”. To ensure sustained attention to the neutral face, despite potential boredom or fatigue, participants were informed they would need to answer detailed questions about that face at the end of the experiment. The emotion words written under the adaptors were never mentioned to the participants.

Before and after the experiment, participants were presented with the mood scale, and at the end of the experiment, participants responded to open-ended questions about the experiment (see questions and responses on: https://osf.io/wk9xq/?view_only=aabe0f707ce243e180c1ed34a8d0bfc7).

### 4.2 Results

The mean PSE in the happy condition (M = 42.19; SD = 16.43) did not differ significantly from the mean PSE in the sad condition (M = 42.39; SD = 12.62), t(20) = −.079, *p* = .938, *d_z_* = −.017 (see Figure 3**: Words**), which shows that the presentation of emotional words alone does not influence participants’ judgments. A BF_10_ = .228 indicated moderate evidence for the null hypothesis of no effect.

#### Mood scores and self-reports

Participants’ mean mood ratings before (M = 63.11; SD = 26.82) and after (M = 66.85; SD = 22.63) the experiment did not differ significantly, *t*(19) = .973, *p* = .343, *d_z_* = .218.

#### Task strategies overview from follow-up questionnaires

Approximately 86% of participants reported focusing primarily on the eyes and other facial features. When asked about their thoughts during the task, about 62% analyzed the emotional expression of the face in reference to the words, with 29% of them noting a discrepancy between the observed face and the emotion word. 90% of participants reported paying attention to the words, although some did so only partially or initially. The perceived influence of these words on participants’ answers was mixed, with 52% reporting being affected by the words, 38% claiming their answers remained unaffected, and the last 10% not being sure or reporting not seeing any words. When asked to guess the experiment’s purpose, 52% expected the words under the adaptors to influence their answers, with only one person guessing assimilative aftereffects. Finally, 57% correctly answered the question that they should judge the test faces (those shown for a shorter period), while 43% made comments unrelated to this question. The exact report of the answers can be found in the Supplementary Materials: https://osf.io/wk9xq/?view_only=aabe0f707ce243e180c1ed34a8d0bfc7.

## 5. Experiment 4

Contrastive aftereffects are typically observed when participants are exposed to physical stimuli depicting extreme emotions; viewing a happy face, for example, generally makes subsequent faces appear sadder (e.g., D’Ascenzo et al., 2014; DeBruine et al., 2010). However, in our previous experiments, participants were not presented with physical extremes but rather engaged in mentally constructed extremes through suggestion (experiment 1a,1b) or imagery (experiment 2). In these experiments, we observed assimilative rather than contrastive aftereffects. To ensure our paradigm’s validity and to confirm that it can elicit both types of aftereffects, we conducted Experiment 4, adapting participants to actual extreme expressions and testing the same stimuli as in our previous experiments.

### 5.1. Materials and methods

#### Participants

N = 23 participants took part in experiment 4. There were no outliers based on the imposed criteria. The full sample (11 females) had a mean age of 24.91 years (SD = 3.49). 21 participants were right-handed and one participant was left-handed (M = 86.52; range: −100 – 100). Participants were tested individually or in groups, each seated in separate cabins.

#### Stimuli

Adapting stimuli consisted of the happy (AF29HAS) and sad (AF29SAS) expressions used in the previous experiments as extremes to construct the continuum of test stimuli (see Figure 2b**: Exp. 4. Perception**).

#### Experimental procedure

The methods and analyses, including the practice session, were identical to Experiment 3 (see Fig. 2), with the only difference being that the adapting stimuli were replaced by an actual happy or sad face. Participants were instructed to maintain their gaze on adaptors. As in Experiment 3, to motivate participants to continuously look at adapting stimuli, they were told that at the end of the experiment they would respond to questions asking details about them.

### 5.2. Results

There was a significant difference between PSEs across the two conditions (see Figure 3**: Perception**): The mean PSE in the happy condition (M = 28.62; SD = 15.54) was significantly lower than the mean PSE in the sad condition (M = 69.90; SD = 13.11, t(22) = −13.122, p < .001), with an extremely large effect size (d*_z_* = −2.736), reflecting contrastive aftereffects, i.e., adaptation to a happy emotion biases the perception of test faces towards sadder emotion, and the opposite for an adaptation to a sad face. A large Bayes factor (BF_10_ = 1.170×10^+9^) provides extreme evidence supporting the contrastive aftereffect.

#### Mood scores

Participants started and finished the experiment with mood ratings of M = 79.27 (SD = 18.35), and M = 74.82 (SD = 18.52), and these ratings did not differ significantly from each other, z = 1.120, p = 0.270, r = 0.273.

#### Task strategies overview from follow-up questionnaires

Across all 23 participants, everyone consistently reported that they looked at the face that stayed on the screen for a longer period. Regarding their thoughts while observing the faces, 61% analyzed the emotional content of the faces, while smaller percentages focused on other aspects, such as remembering features (9%), or had experiences like confusion or frustration (4%). When judging happy or sad faces, 65% of participants reported judging the test faces, 13% indicated judging the adaptors, and 22% provided answers unrelated to the question. In terms of understanding the experimental manipulation, 17% showed awareness of contrastive adaptation effects, 9% made guesses about assimilative effects, 30% believed the adaptor influenced their judgments but did not specify how, and 26% seemed unclear on the experimental manipulation. For the exact responses refer to: https://osf.io/wk9xq/?view_only=aabe0f707ce243e180c1ed34a8d0bfc7.

## 6. General Discussion

This study examines whether two cognitive strategies, mental imagery and autosuggestion, impact the perception of facial emotions. In an adaptation paradigm, participants were asked to use autosuggestion to *perceive* the expression of a neutral female face as happier or sadder than it actually is (Experiment 1a and 1b) or to *imagine* the previously seen neutral face as being happy or sad (Experiment 2). In all three experiments, participants judged test faces as happier or sadder, following the happy and sad adaptations, respectively, both when performing autosuggestion or when imagining the respective emotion. Critically, we found no effects of adaptation to facial neutral expressions that were accompanied by emotional words (Experiment 3), suggesting, that the effects observed in the first three experiments were not an expression of response bias, but the results of a cognitive process centered on mental imagery and autosuggestion, respectively. Finally, we observed classical, contrastive aftereffects when a happy or a sad face was used as adaptor (Experiment 4), further supporting the notion that our paradigm successfully replicated established perceptual adaptation aftereffects (D’Ascenzo et al., 2014; DeBruine et al., 2010; Ellamil et al., 2008; Kaping et al., 2002). Together, our experiments show that both autosuggestion and mental imagery can be used to modulate emotion perception in a social context.

Experiment 1a showed that autosuggestion modulates emotion perception in such a way, that the emotions in test faces resemble more the adapting faces. However, as most participants of experiment 1a reported creating visual images to support their autosuggestion, despite instructions to avoid imagery, experiment 1a could not fully separate the processes of imagery versus autosuggestion. In experiment 1b, therefore, revised instructions emphasized verbal suggestion, reducing reliance on imagery. This led indeed to an increased reliance on verbal suggestions and a decrease in imagery use. However, participants still often combined verbal strategies with mental imagery and other techniques. The tendency to use imagery in the autosuggestion group highlights several points: participants may conflate autosuggestion with imagery despite understanding the distinction; they might simply lack the skills to effectively use autosuggestion in the form of intentional thought repetition without associated imagery; or individuals with strong imagery abilities may find it difficult to avoid using them, naturally combining both techniques to enhance outcomes. Autosuggestion appears to be a flexible process that engages multiple cognitive strategies, and the extent to which participants rely on imagery or verbal repetition may depend significantly on task instructions and individual tendencies. Future research could explore autosuggestion in individuals who are unable to create mental images (aphantasics, e.g., Monzel et al., 2023; Reeder et al., 2024) to help distinguish pure thought repetition from imagery-supported autosuggestion.

Nonetheless, despite the strong interplay between these processes, the tasks likely differ in their cognitive demands and underlying intentions. In the “imagery only” task, participants create an emotional face in the absence of any visible stimulus. In contrast, the autosuggestion task involves actively altering the perceived emotion of a physically present stimulus by reinterpreting a neutral face as happy or sad. This distinction holds significant implications for daily experiences, suggesting the possibility of intentionally altering one’s perception of another’s facial expression, especially in face-to-face interactions.

Experiments 1a-b and 2 resulted in assimilative aftereffects, whereas experiment 4, which used physical extreme stimuli, resulted in contrastive aftereffects. One possible interpretation of these differing effects aligns with the findings by Korolkova (2018), who noted a distinction in the direction of aftereffects based on the nature of the adapting stimuli. Specifically, the study found contrastive aftereffects with static images of emotional faces, whereas assimilative aftereffects were observed with stimuli that dynamically alternated between expressions, such as disgust and happiness. Our findings might parallel this distinction. In Experiments 1a-b and 2, participants reported placing the imagined or autosuggested emotion in a situational context. In the imagery condition, they generated mental representations of scenes rather than holding one static image of a face in the mind’s eye. Similarly, participants feedback indicated that when autosuggesting a specific emotion, the image they looked at on the screen, fluctuated in perceived intensity of the suggested emotion, aligning with the dynamic qualities associated with assimilative effects proposed by Korolkova (2018).

The assimilative effects observed may also reflect priming on participants’ judgments. For example, Carroll and Young (2005) demonstrated that emotional priming, using words or emotion-related stimuli presented for only 0.5 s immediately before the test stimuli, led to faster and more accurate emotion recognition when the primes matched the target facial expressions. In a complementary approach, Fischer and Whitney (2014) proposed that perception is dependent on both prior and current perceptual information, a phenomenon termed *serial dependence*, which fosters continuity in perception despite environmental noise. To illustrate this point, Liberman and colleagues (2014) showed that participants’ perception of facial identity was biased towards identities seen up to several seconds before, independent of their previous responses. Notably, the 7-second time difference between viewing a previous face and a current target face, as reported by Liberman and colleagues, corresponds to the delay between the onset of the imagined/autosuggested face and the presentation time of the test face in our paradigm (see Figure 2a), which may suggest a tendency toward perceptual continuity in our findings.

Contrastive aftereffects are commonly observed with perceptual stimuli, as demonstrated in Experiment 4, where participants are typically exposed to adaptors representing an extreme end of the stimulus spectrum, also in the context of high-level adaptation aftereffects (Ambroziak et al., 2023; Clifford & Rhodes, 2005; Myga et al., 2024a; Pell & Richards, 2011). Although one might speculate that the milder, mentally generated adaptors in Experiments 1a-b and 2 could account for the directional differences observed, this is unlikely, as the intensity of adapting stimuli generally influences aftereffect strength rather than direction (Burton et al., 2013; Calder et al., 2008; Hong & Yoon, 2017; Lawson et al., 2009; Rhodes et al., 2017).

While multiple interpretations for the aftereffects are plausible, the primary aim of this study was not to differentiate whether the observed effects stem from adaptation, priming, or serial dependence, but rather to assess our approach’s efficacy in influencing subsequent perceptions and determining the effect’s direction. To address potential response bias (Nichols & Maner, 2008; Orne, 1962), we conducted Experiment 3, exposing participants to neutral faces together with “HAPPY” or “SAD” labels. Contrary to demand characteristics or cognitive expectations, participants showed no consistent bias based with the labels. Although many participants noticed the words and believed they should influence their judgments, our findings indicate that the deliberate intent to perceive specific emotions, as in Experiments 1a-b and 2, is necessary for significant shifts in emotion perception. Experiment 3 thus supports the idea that the effects observed in the earlier experiments extend beyond simple response bias and highlight the role of active and deliberate cognitive engagement in altering facial emotion perception.

As an exploratory measure, we asked participants to rate their mood as well as to indicate more details on how they evoked imagery or autosuggested states in themselves. Evidence suggests that interactions between self-mood and emotion processing may be significant (Sel et al., 2015). In Experiment 1b, participants experienced a mean mood drop of 9 units after autosuggesting a sad emotion, though this change was not statistically significant. This within-condition comparison was exclusive to Experiment 1b, as the other experiments included mood ratings solely at the beginning and end. Additionally, in both the autosuggestion and imagery experiments, the majority of participants reported modifying their facial expressions or emotional states to facilitate the generation of mental images and autosuggestive outcomes. This finding aligns with research demonstrating that modulating facial mimicry – either by inhibiting or enhancing it – can influence the recognition of emotional facial expressions (Rychlowska et al., 2014; Sel et al., 2015).

Our results might be an early indicator that exposure to certain emotional stimuli could potentially be exploited as a tool to bias emotion perception in the longer term. However, while the endurance of adaptation aftereffects in various aspects of face perception have been extensively studied, including those related to identity (Carbon et al., 2007; Carbon & Ditye, 2011), investigations into long-lasting adaptation effects in emotion perception seem lacking. Carbon and Ditye (2011) conducted a series of experiments where participants exhibited sustained adaptation aftereffects, lasting up to a week, following exposure to distorted images of familiar faces. These effects not only persisted over time but also generalized across different images of the same individual and even extended to other identities. Along these lines, it has been proposed that body shape misperception, as in eating disorders, might be an example of long-term adaptation aftereffects caused by recalibration of the perceptual system to the exposure to a constant media extreme of “ideal”, very thin, bodies (Brooks et al., 2020). In that regard, exposure to bodies of larger size might serve as a therapeutic intervention in patients with eating disorders (Challinor et al., 2017; Porras-Garcia et al., 2020).

In light of these findings, our study might similarly recalibrate emotional perception, potentially biasing the interpretation of others’ facial expressions. Future research should explore the durability of these effects in emotion perception and assess whether prolonged exposure to specific perceptual, imagined or suggested emotional stimuli could lead to enduring perceptual biases. Nonetheless, several limitations of this study need to be considered. First, we used the same facial identity throughout the study, limiting the generalizability of our findings, and the sample primarily consisted of individuals from or residing in Europe, which may restrict broader applicability. Additionally, while our effect sizes were substantial, with moderate Bayes factors for imagery and large Bayes factors for autosuggestion, they were not as large as those observed with physical stimuli, and the effects were not uniform across all participants, suggesting limitations in the potential practical application of autosuggestion and imagery techniques. Furthermore, despite efforts to control for response bias, it remains possible that such biases contribute to the observed assimilative aftereffects, a limitation that is inherent to all studies showing assimilative aftereffects in the literature. Another important consideration is whether our procedures genuinely bias perception, influence the interpretation of emotional stimuli, or both. Additionally, we cannot definitively conclude whether autosuggestion and mental imagery represent distinct tasks, as the first three experiments might have tapped into the same underlying mechanism, only differing in the presence or absence of the neutral physical stimulus. Although we believe that participants approached each task with different intentions, and therefore engaged in different cognitive processes, we cannot be certain of this distinction.

Finally, we included various self-report questionnaires to capture individual differences in task performance. While our analysis was broad, focusing on general trends rather than on individual differences, the self-reports offered important insights into how participants approached the tasks. For instance, many participants in Experiment 1a omitted the verbal repetition component of autosuggestion, focusing instead on visualizing the suggested emotion. Similarly, in the mental imagery condition, participants often did not evoke the identity of the neutral face but engaged in other forms of visualization based on their emotional targets. These findings underscore the importance of understanding participants’ interpretations of task instructions and their strategies for performing cognitive tasks. Indeed, overlooking this information, as is often the case in cognitive neuroscience, prevents a full understanding of whether the cognitive process intended by the study design is indeed the one participants engage in.

Concluding, our study contributes with evidence that exposure to emotional stimuli through autosuggestion or imagery can lead to a consistent bias in perceiving or interpreting recognizing facial emotions in other people. These results highlight the potential for training and deliberate application of self-directed interventions, such as imaginative and autosuggestive techniques, to shape perception of emotional expression.

## Data Availability

Testing took place in 2024 in Magdeburg, at the Leibniz Institute for Neurobiology. Materials and data are available on the project OSF page: https://osf.io/wk9xq/?view_only=aabe0f707ce243e180c1ed34a8d0bfc7. This includes all raw, and processed data, Matlab scripts, instructions given to participants, and written reports of strategies used by participants.

## Acknowledgments

We would like to thank Brinda Valsaraj for helping with participant recruitment for experiments 1a and 2.

## Declaration of Generative AI and AI-assisted technologies in the writing process

During the preparation of this article, the authors used ChatGPT to improve the readability and language of the manuscript. After using this tool, the authors reviewed and edited the content as needed and take full responsibility for the content of the published article.

## Author contributions

- Conceptualization: KAM, EA
- Data curation: KAM, EA
- Formal analysis: KAM, EA
- Funding acquisition: KAM, EK, EA
- Investigation: KAM, MRL, EK, EA
- Methodology: KAM, EA
- Project administration: KAM, EA
- Resources: EA
- Software: EA
- Supervision: EA
- Validation: KAM, EA
- Visualization: KAM
- Writing original draft: KAM
- Writing: review and editing: KAM, MRL, EK, EA

